# Forward processing of spatiotemporal maps across hippocampal subfields in common marmosets

**DOI:** 10.64898/2026.04.28.721285

**Authors:** Feng Qiao, Yoshio Sakurai, Atsushi Nambu, Anna Roe Wang, Yuji Naya

## Abstract

Episodic memory requires distinguishing similar events occurring at the same location but at different times. While the hippocampus integrates “where” and “when,” the specific roles of the primate dentate gyrus (DG) and CA3 remain elusive. We recorded single-unit activity from freely moving marmosets navigating a spatial maze in a visually impoverished environment. Both hippocampal subfields robustly encoded self-position via neurons selective for stationary epochs at platforms and movement along trajectories, whereas gaze-related coding was minimal. While DG signaled stationary and dynamic self-positions independently, CA3 integrated these elements into conjunctive representations. Critically, only CA3 neurons exhibited strong temporal-order modulation. These findings reveal a feedforward disambiguation process in which CA3 transforms discrete spatial inputs from DG into the integrated spatiotemporal maps essential for episodic memory.

## Introduction

Distinguishing between highly similar experiences—such as meeting the same friend at the same coffee shop on different days—requires assigning each event to a specific temporal context. This disambiguation of overlapping spatial episodes by their timing is a hallmark of episodic memory (*1*). The hippocampus is central to this process, providing a scaffold that integrates “where” and “when” information into distinct spatiotemporal memory traces (*2, 3*). However, how the differential processing of these components by hippocampal subfields in primates is still under debate. To keep similar events attached to distinct spatiotemporal contexts, the hippocampus is thought to perform pattern separation. Classic computational accounts, notably Marr’s theory of the archicortex (*4*), posit a functional hierarchy: dentate gyrus (DG) decorrelates incoming signals through expansion recoding, whereas CA3, with its dense recurrent collaterals, supports auto-association and constructs integrated representations (*5*). Rodent studies support roles for both DG and CA3 in pattern separation of spatial contexts (*6*), but it remains unclear how these subfields contribute to integrating spatial location and temporal order to separate overlapping episodes. The origin of temporal signals that modulate spatial representations is also unresolved. “Time cells” have been robustly identified in CA1 (*7*), yet it is unclear whether temporal coding emerges within DG and CA3 or is inherited from downstream fields or cortical feedback. Clarifying how temporal-order information—such as distinguishing the first from the third visit to the same location—is incorporated into hippocampal circuits is crucial for understanding how the brain disambiguates sequential task phases.

Rodent work has traditionally focused on “place cells” that encode physical self-position (*8*). In contrast, primate studies have emphasized “spatial view cells” that fire according to gaze location in allocentric coordinates, even with fixed body position (*9–12*). Here, we address these questions using the common marmoset, a primate model well suited for studying free navigation and vision. Using a visually guided travel (VGT) task in a spatial maze, we asked whether hippocampal neurons in marmosets primarily code self-position or spatial view in a visually constrained environment, and how temporal-order information is integrated with these spatial representations. By recording single-unit activity from DG and CA3 during the VGT task, we reveal a hierarchical organization of episodic information processing in the primate hippocampus and provide evidence for forward processing of spatiotemporal representations along the DG–CA3 axis.

## Results

### Spatial maze and visually guided travel task

Two common marmosets were trained to perform a VGT task in a two-dimensional spatial maze (Fig. 1A, S1A). The maze consisted of a black floor (2.0 m ×1.5 m) surrounded by four black walls (0.7 m height). The maze was covered by a transparent plastic roof and placed in a dark room without visible external landmarks. Four white circular platforms (10 cm in diameter and 1.0 cm high), each with an LED attached underneath, were located near walls. Among the four platforms, three test platforms (Platforms *A*, *B*, and *C*) were arranged in an equilateral triangle (with a 1.1 m distance between each platform), and the home platform (Platform *H*) was located adjacent to the reward sipper. Head position and direction were monitored via LEDs mounted on the animals’ heads.

**Fig. 1.**
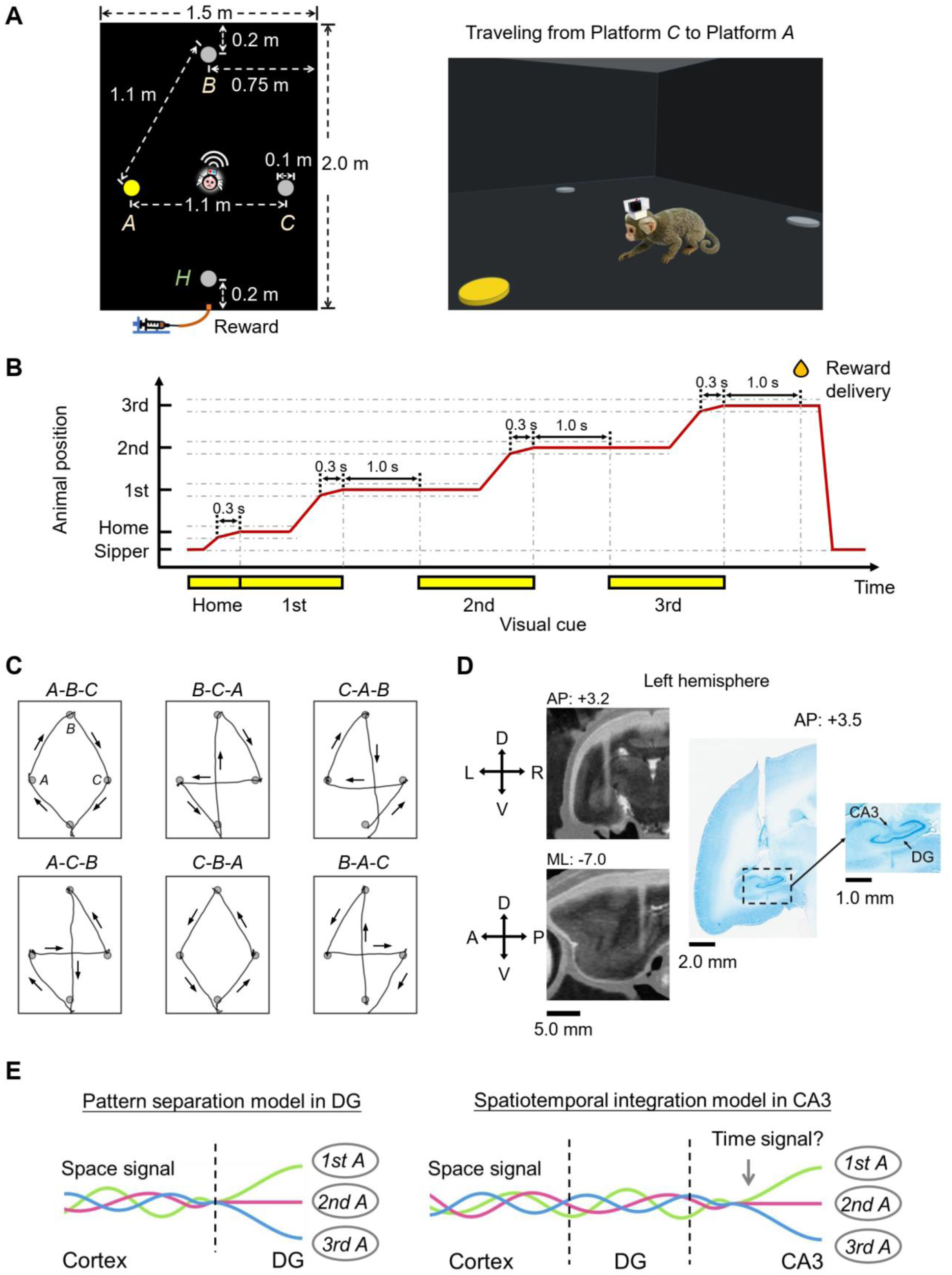
Experimental setup and task design of the visually-guided travel (VGT) task. (A) Schematic illustration of the arena and platform configuration. Top-view (left) and 3D perspectives (right) of the maze. The arena floor was black, with gray circles representing platforms and yellow indicating platforms with activated LEDs. *H* denotes the home platform, and *A*, *B*, *C* represent test platforms, respectively. A reward-sipper was positioned adjacent to the home platform (*H*), at a height of 10 cm above the floor. (B) Timeline depicting the temporal sequence. The diagram illustrates the activation of visual cues, marmoset movement, and reward delivery in a single VGT trial. Yellow bands represent LED illumination beneath the 1st, 2nd, and 3rd target platforms. The red line indicates the marmoset’s movement trajectory, reflecting its positional dynamics throughout the trial. Upon entering the current target platform zone, the animal was required to maintain its position for 1.3 s to trigger illumination of the next target LED. A reward was administered following successful completion of all three sequential traveling steps. (C) Six sequential patterns employed in the VGT task. Gray circles represent platforms, black lines denote the marmoset’s trajectory, and arrows indicate the direction of locomotion. (D) Electrode localization. Final electrode positioning relative to hippocampal subregions for Monkey B. (Left) MRI and CT imaging showing the electrode track. (Right) Nissl-stained coronal section at the target level, displaying the electrode track relative to the dentate gyrus (DG) and CA3 subregions of the hippocampus. (E) Proposed computational model. A comparison of the pattern separation model in the DG (left) and the spatiotemporal integration model in the CA3. Twisted tri-color strands represent spatial signal encoding self-position at Platform *A*, which remain indistinguishable across temporal orders in the DG. Conversely, untwisted strands in the CA3 illustrate the distinct spatiotemporal representation representations that encode both “when” and “where” information.

In the VGT task, animals were required to travel over one home and three test platforms according to the illumination of LEDs underneath them (Fig. 1B; Movies S1-S3). Each trial began when the animal stayed at Platform *H*. The animal then had to visit one of three test platforms indicated by LED illumination and stay there at least 1.3 s. Then, another platform was illuminated. After the animal had visited all three test platforms, a reward (liquid food) was delivered through the sipper. Six travel patterns (*A*-*B*-*C*, *B*-*C*-*A*, *C*-*A*-*B*, *A*-*C*-*B*, *C*-*B*-*A*, and *B*-*A*-*C*) were presented pseudo-randomly across trials (Fig. 1C). During the recording phase, Monkey B completed 32.6 ±6.0 (mean ±s.d.) trials per session, and Monkey F completed 28.0 ±3.1 trials per session (Table S1).

Neural signals were acquired via chronically implanted 16-channel brush-array electrodes mounted on micromanipulators and recorded wirelessly (Fig. S1B, S1C); recording sites were verified using X-ray CT, MRI, and histological reconstructions (Fig. 1D, S1D). We recorded single-unit activity from CA3 (224 neurons from 40 sessions) and DG (143 neurons from 22 sessions) while the animals performed the VGT task.

### Self-position representation during stationary epochs at platforms

We first examined how self-position influenced neuronal activity while animals were stationary on the three test platforms. Figure 2A illustrates a DG neuron that fired robustly when the animal stayed on Platform *B* (5.10 ±0.95 Hz, mean ±SEM) but showed almost no activity on Platforms *A* (0.05 ±0.03 Hz) or *C* (0.08 ±0.04 Hz), reflecting strong selectivity for self-position across the platforms (*P* < 0.0001, *F* [2, 61] = 35.9, one-way ANOVA). We defined “platform neurons” as neurons that exhibited significantly different firing rates across the three test platforms (*P* < 0.01, one-way ANOVA). The proportion of platform neurons greatly exceeded the 1% expected by chance in both CA3 (35.3%; *P* < 0.0001, *Z* = 51.5, binomial test) and DG (29.4%; *P* < 0.0001, *Z* = 34.1; Table 1), and did not differ significantly between the two subregions (*P* = 0.24, *χ*^2^ = 1.37, *χ*^2^ test).

**Fig. 2.**
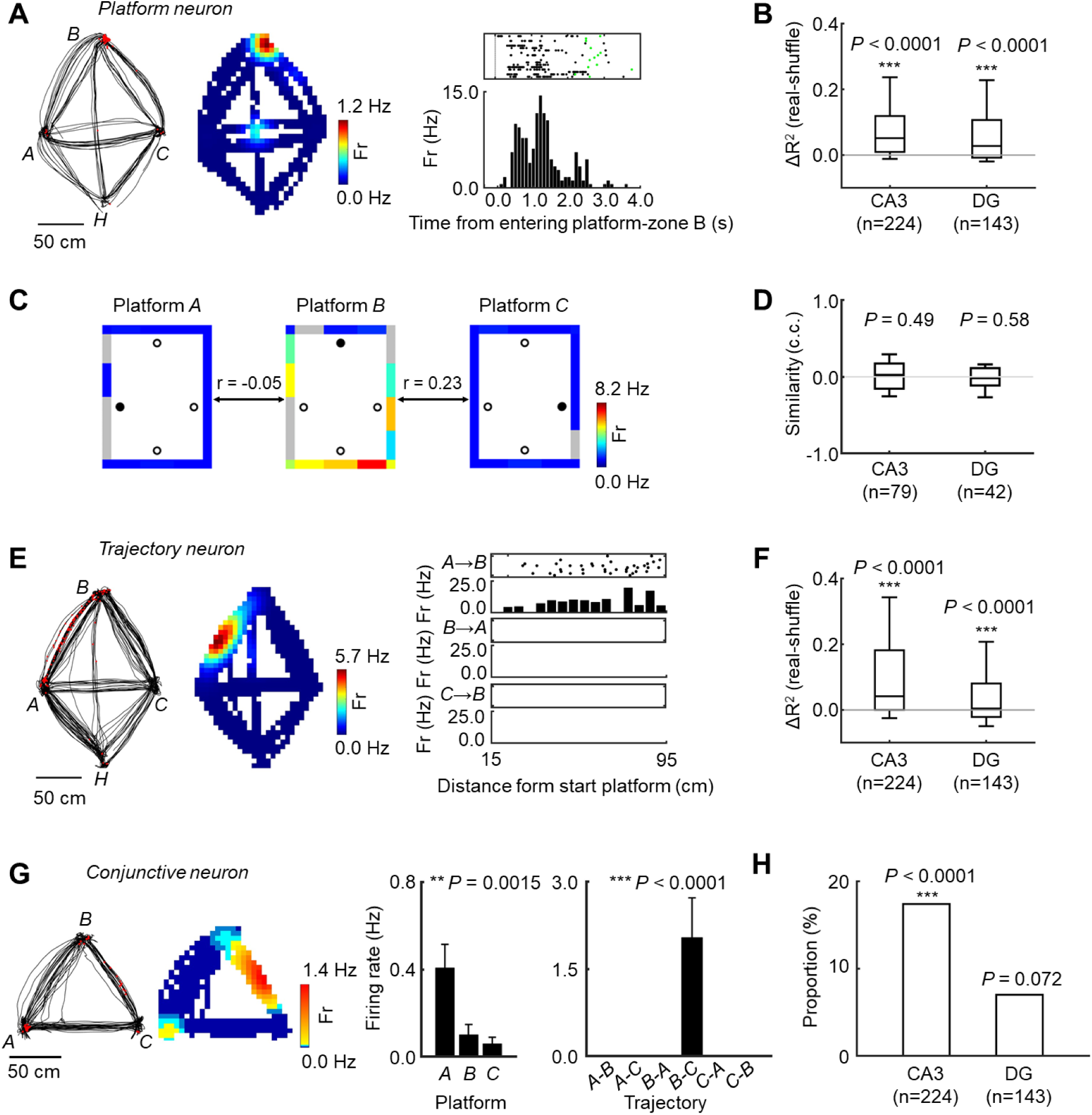
Self-position selective activities. (A) Example platform neuron in DG of monkey F. (Left) Response map during the VGT task. Black lines represent trajectories, and red dots indicate spikes. Color represents the firing rate at each location. (Right) Raster displays aligned to the time the marmoset entered Platform *B* zone. Green dots represent time points when the marmoset exited the zone. (B) Distributions of *R*^2^ values reflecting platform-selective activity. Box-and-whisker plots for all recorded neurons in CA3 and DG. Each *R*²value was normalized by subtracting the mean of 1,000 shuffled *R*²values for that neuron. The horizontal line within each box represents the median; box boundaries indicate the interquartile range (IQR). Whiskers extend to the 10th and 90th percentiles. (C) Facing-location analysis for the example platform neuron. The response pattern is displayed according to the animal’s facing location at each stationary test platform (black filled circles). Colored strips along the wall represent mean firing rates for each 50-cm spatial bin, corresponding to the animal’s facing location toward the maze wall. Gray bins were excluded due to insufficient occupancy (< 1.0 s). *r* denotes the Pearson correlation coefficient reflecting firing pattern similarities. (D) Firing pattern similarities for facing location. Pearson correlations were calculated between the best-preferred platform and the other two platforms; the mean of these two similarities was calculated for each platform neuron. Box-and-whisker plots indicate the distributions of these mean similarities in CA3 and DG. (E) Example trajectory neuron in the CA3. (Left) Response map in the same format as Fig 2A. (Right) Raster displays during travel along Trajectories *A*→*B*, *B*→*A* and *C*→*B*. The bar graph indicates mean firing rates at each spatial bin (5 cm) along the trajectory. (F) Distributions of *R*² values reflecting trajectory-selective activity. Box-and-whisker plots for all recorded neurons in CA3 and DG. *R*²values were normalized by subtracting the mean of 1,000 shuffled *R*²values for each neuron. (G) Example conjunctive neuron. (Left) A CA3 neuron exhibiting both platform-selective and trajectory-selective responses. (Right) Bar graphs indicate mean firing rates (± SEM). (H) Proportions of conjunctive neurons among all recorded neurons in CA3 and DG.

**Table 1.** Numbers of neurons coding self-position information.

|  | **Recorded** | **Self-position** | **Platform** | **Trajectory** |
| --- | --- | --- | --- | --- |
| **Total** |  |  |  |  |
| **CA3** | 224 | 104 (46.4 %) | 79 (35.3 %) | 64 (28.6 %) |
| **DG** | 143 | 54 (37.8 %) | 42 (29.4 %) | 22 (15.4 %) |
| **Monkey B** |  |  |  |  |
| **CA3** | 151 | 73 (48.3 %) | 56 (37.1 %) | 49 (32.5 %) |
| **DG** | 46 | 14 (30.4 %) | 11 (23.9 %) | 5 (10.9 %) |
| **Monkey F** |  |  |  |  |
| **CA3** | 73 | 31 (42.5 %) | 23 (31.5 %) | 15 (20.6 %) |
| **DG** | 97 | 40 (41.2 %) | 31 (32.0 %) | 17 (17.5 %) |

Platform and trajectory neurons exhibited significant platform-selective and trajectory-selective activity, respectively ( $P < 0.01$ for each, one-way ANOVA). “Self-position neurons” are defined as those exhibiting either significant platform-selective or trajectory-selective activity.
Percentages of parentheses indicate proportions of significant neurons among recorded neurons.

To quantify the strength of self-position coding across the entire population, we computed an *R²*value for each neuron, representing the proportion of firing-rate variance explained by platform identity. Observed *R*^2^ values of the recorded neurons were significantly higher than their respective means of 1,000 shuffled *R*²values in both CA3 (*P* < 0.0001, n = 224, *Z* = 11.2, Wilcoxon signed-rank test, two-tailed) and DG (*P* < 0.0001, n = 143, *Z* = 6.75) (Fig. 2B), consistent with the elevated proportions of platform neurons. Moreover, the normalized *R*²values by subtracting the shuffle means were significantly larger in CA3 than in DG (*P* = 0.015, *KS* = 0.166, Kolmogorov–Smirnov test; Fig. S2A), indicating somewhat stronger self-position coding in CA3 at platforms although tuning properties such as contrast and sharpness of platform neurons did not differ between the two subregions (Supplementary Text, Fig. S2B–D).

### Self-position coding is not explained by facing direction or facing location

During stationary periods on the test platforms, animals showed a strong bias in their facing directions (Fig. S3A). For example, when the animals stayed at Platform *C*, they tended to direct their faces to the reward sipper (Fig. S3B). Because this bias produced different patterns of facing location across platforms, a potential concern is that “platform selectivity” might reflect differences in facing direction or facing location rather than self-position per se. For example, the strong response of the example neuron at Platform *B* could, in principle, be attributed to the locations toward which the animal was facing while at Platform *B*.

To disentangle self-position from facing location, we projected the animal’s facing direction at each test platform onto the maze walls and computed mean firing rates in 50-cm bins along the wall (up to 14 bins), considering only bins with > 1.0 s of total occupancy across trials. The representative neuron shown in Fig. 2A fired strongly when the animal was at Platform *B* facing toward Platform *C* and the reward sipper, but exhibited little or no activity when the animal faced the same locations from Platforms *A* or *C* (Fig. 2C). To assess whether responses at the preferred platform could be explained by facing location, we calculated Pearson correlation coefficients between the firing-rate distributions across facing locations at the preferred platform and each of the two non-preferred platforms. For the example neuron, these correlations were not significant for either Platform *A* (*r* = −0.05, *P* = 0.90, *d.f.* = 7) or Platform *C* (*r* = 0.23, *P* = 0.55, *d.f.* = 7), confirming that the neuron encoded self-position rather than facing location.

Across platform neurons, the mean correlation between facing-location firing-rate maps at the preferred versus the other two non-preferred platforms did not differ from zero in either CA3 (median = 0.022, *P* = 0.49, n = 79, *Z* = 0.690; Wilcoxon signed-rank test, 2-tailed) or DG (median = −0.015, *P* = 0.58, n = 42, *Z* = -0.556; Fig. 2D). Extending this analysis to all recorded neurons, using all three possible platform pairings, we again found no significant positive correlations in either region (CA3: median = 0.000, *P* = 0.43, n = 224, *Z* = 0.793; DG: median = −0.022, *P* = 0.51, n = 143, *Z* = -0.664; Fig. S3C). These findings indicate that platform-selective activity in both CA3 and DG primarily encodes self-position, rather than facing location, during the VGT task.

We also assessed the overall influence of facing direction on neuronal spiking, independent of task events, using a Rayleigh test (*13*). Only a negligible fraction of neurons showed significant modulation by facing direction in either region (CA3: 0.45%, *Z* = −0.833, *P* = 0.89; DG: 0.70%, *Z* = −0.361, *P* = 0.76; binomial test). These results further support the conclusion that hippocampal activity while at platforms predominantly reflects self-position rather than head direction at least in the visually impoverished environment.

### Space representation during travel between platforms

We next examined the effect of the animals’ self-position while traveling from one test platform to another. Fig. 2E shows a CA3 neuron that exhibited a strong response when the animal traveled along the trajectory from Platform *A* to Platform *B* (Trajectory *A*→*B*, 7.7 ± 1.8 Hz), but no response for other trajectories, including the opposite direction (Trajectory *B*→*A*) and a trajectory with the same goal destination (Trajectory *C*→*B*). A one-way ANOVA with six trajectories as the main factor revealed a significant trajectory-selective response in this neuron (*F* [5,52] = 16.3, *P* < 0.0001). We defined neurons with a significant (*P* < 0.01) trajectory effect as “trajectory neurons.” The proportion of trajectory neurons was significantly higher than 1% expected by chance in both CA3 (28.6%; *P* < 0.0001, *Z* = 41.5; binomial test) and DG (15.4%; *P* < 0.0001, *Z* = 17.3) although the proportion of trajectory neurons was larger in CA3 than in DG (*χ*^2^ = 8.46, *P* = 0.0036; *χ*^2^ test; Table 1). The substantial number of trajectory neurons in both subfields was further supported by *R*²values, which estimated the effect of trajectory compared with means of 1,000 shuffled *R*²values (CA3: n = 224, *Z* = 8.93, *P* < 0.0001; DG: n = 143, *Z* = 4.08, *P* < 0.0001; Wilcoxon signed-rank test, two-tailed; Fig. 2F). Consistent with the proportion of trajectory neurons, the *R*²values normalized by subtracting the shuffle means were significantly larger in CA3 than in DG (*KS* = 0.16, *P* = 0.024; Kolmogorov-Smirnov test; Fig. S4A), although the tuning properties of trajectory neurons, evidenced by their exclusive responses to the best-preferred trajectory, were similar in both subfields (Supplementary text, Figs S4B-E).

The exclusive coding of the best-preferred trajectory suggests direction-selective responses in trajectory neurons (Fig. 2E). However, trajectories sharing the same spatial positions might induce correlated responses between trajectory pairs bound to the same platforms (e.g., Trajectory *A*→*B* and Trajectory *B*→*A*). To test this, we calculated Pearson correlation coefficients between responses to these trajectories pairs. The correlation coefficients did not differ significantly from zero in either subregion (CA3: median = 0.0, *P* = 0.46, *Z* = -0.735; DG: median = 0.0, *P* = 0.44, *Z* = -0.776; Wilcoxon signed-rank test, two-tailed; Fig. S4F, “Position”). We also assesed whether trajectory neurons exhibited correlated responses to trajectory pairs toward the same goal platform (e.g., Trajectory *A*→*B* and Trajectory *C*→*B*). Again, trajectory-selective responses could not be explained by the goal location toward which the animal was facing during travel (CA3: median = 0.0, *P* = 0.089, *Z* = 1.70; DG: median = −0.32, *P* = 0.062, *Z* = -1.87; Wilcoxon signed-rank test, 2-tailed; Fig. S3E, “Goal”). Thus, although trajectory neurons showed strong direction selectivity, their responses did not reflect a specific facing location, consistent with the properties of platform neurons.

### Space representation at stationary platforms and traveling trajectories

During the VGT task, a substantial fraction of neurons signaled the animals’ self-position, either while stationary on platforms or while traveling between platforms, in both CA3 (46.4% of recorded neurons) and DG (37.8%) (Table 1). Among these “self-position neurons” (platform- or trajectory-selective), some exhibited both platform-selective activity and trajectory-selective activity. Figure 2G presents an example CA3 neuron with a conjunctive spatial representation coding Platform *A* and Trajectory *B*→*C*. The proportion of these conjunctive neurons was significantly higher in CA3 (17.4 % of recorded neurons) than in DG (7.0 %) (*P* = 0.0042, *χ*^2^ = 8.19) (Fig. 2H). Moreover, chi-square tests examining an independency between platform-selective activity and trajectory-selective activity revealed that CA3 neurons tended to exhibit both types of activity, whereas DG neurons did not (*P* < 0.0001, *χ*^2^ = 25.9) but not in DG (*P* = 0.072, *χ*^2^ = 3.24). These results suggest that CA3 integrates spatial information regarding stationary platforms and traveling trajectories into conjunctive representations, whereas DG neurons tend to represent these two types of spatial information independently.

### Temporal-order effects on spatial representation

We examined temporal-order effects on neural activity while the animals stayed at each test platform during the VGT task. Figure 3A shows an example CA3 neuron whose preferred response to Platform *A* was strongest when the animal stayed there during the first temporal order. To quantify these effects, we applied a two-way ANOVA with platform (‘space,’ three levels) and temporal order (‘time,’ three levels) as main factors. The example neuron showed a significant main effect of tim*e* (*P* < 0.0001, *F* [2,55] = 12.0) and a significant time ×space interaction (*P* < 0.0001, *F* [2,55] = 11.8). We defined neurons with a significant main effect of time or a significant time ×space interaction as “temporal-order neurons.” We found a substantial proportion of temporal-order neurons in CA3 (19.6%, 44/224 recorded neurons; *P* < 0.0001, *Z* = 18.9, binomial test), but not in DG (2.1%, 3/143; *P* = 0.55, *Z* = 0.0840; Table S2). Figures 3B and 3C show CA3 neurons exhibiting the strongest response during the second and third temporal orders, respectively. The dominance of temporal-order effects in CA3 was confirmed by *R*²values for the main effect of time and time ×space interaction: in CA3 (n = 224), both *R*²values were significantly greater than the means of shuffled 1,000 shuffled *R*² values (time effect: *P* = 0.004, *Z* = 2.85; time ×space interaction: *P* < 0.0001, *Z*-value = 5.64; Wilcoxon signed-rank test, two-tailed; Fig 3D). In contrast, this was not observed in DG (n = 143; time effect: *P* = 0.82, *Z* = -0.234; time ×space interaction: *P* = 0.91, *Z* = 0.114). Furthermore, both *R*²values, normalized by subtracting the shuffle means, were significantly larger in CA3 than in DG (*P* = 0.0056, *KS* = 0.181 and *P* = 0.0059, *KS* = 0.180, respectively; Kolmogorov–Smirnov test). These results suggest that temporal-order effects emerge prominently in CA3, which is located one synapse downstream of the DG.

**Fig. 3.**
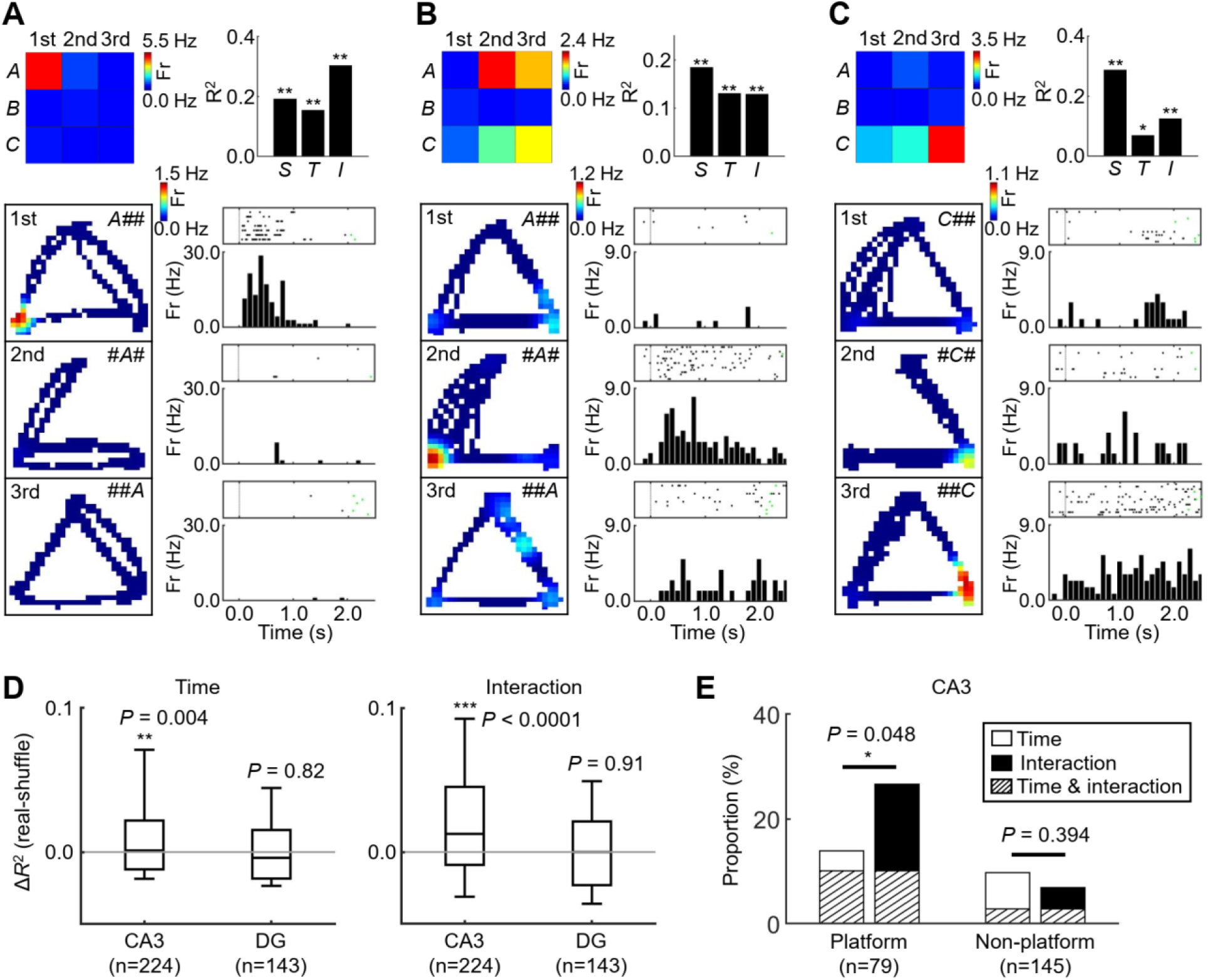
Integration of the space and time information. (A-C) Three example CA3 neurons showing temporal-order effects on platform selective activity. (Top) Color maps indicating firing rates across three different test platforms (rows) and three different temporal orders (columns). Bar graphs indicate *R*^2^ values reflecting main effects of space (*S*) and time (*T*), as well as their interactions (*I*), for each the example neuron (* *P* < 0.05; ** *P* < 0.01; permutation test, n = 1000 shuffles, two-tailed). (Bottom) Spatial firing rate maps when the marmoset was at the best-preferred platform for each temporal order. “*X##”*, “#*X#”,* and “##*X”* denote trials where Platform *X* was the 1st, 2nd and 3rd target, respectively. Raster plots and bar graphs illustrate spike firings and firing rates aligned to the time the marmoset entered the target platform zone. (D) Distributions of *R*^2^ values reflecting temporal modulation. Box-and-whisker plots show the distribution of *R*^2^ values for the main effect of time (left) and the space ×time interaction (right) for all recorded neurons in CA3 and DG. Each *R*^2^ value was normalized by subtracting the mean of 1,000 shuffled *R*^2^ values for that neuron. The format is the same as in Fig. 2B. (E) Proportions of temporal-order neurons. Classification of neurons by the main effect of time and the space × time interaction among platform neurons (left) and non-platform neurons (right).

Next, we examined the relationship between temporal-order effects and self-position coding in CA3. The proportion of temporal-order neurons was significantly higher among platform neurons (30.4%, n = 24/79) than among non-platform neurons (13.8%, n = 20/145; *P* = 0.003, *χ*^2^ = 8.91, *χ*^2^ test). Moreover, CA3 platform neurons (n = 79) showed a time ×space interaction (26.6%, n = 21) more frequently than a main effect of time (13.9%, n = 11) (*P* = 0.048, *χ*^2^ = 3.91, *χ*^2^ test) (Fig. 3E, see also Fig. S5). In contrast, other CA3 non-platform neurons (n = 145) showed a significant interaction (6.9%, n = 10) less frequently than a main effect of time (9.7%, n = 14) although the difference was not statistically significant (*P* = 0.394, *χ*^2^ = 0.727, *χ*^2^ test). These results suggest that CA3 integrates time and space signals into specific combinations representing a particular platform at a particular temporal order, rather than receiving these signals independently.

We also examined temporal-order effects during travel between platforms (Fig. S6A). Although data acquisition was more limited here (approximately one-fifth, Table S1), the results were consistent with those for stationary platform responses. The proportion of temporal-order neurons for traveling trajectories exceeded the chance level (2.0%) only in CA3 (7.6%, n = 17/224; *P* < 0.0001, *Z* = 5.98, binomial test; Table S3). Most of these neurons (82.4%, n=14/17) were also trajectory-selective, and the interaction effect tended to be stronger than the main effect of time (Figs. S6B and S6C), indicating the integration of time and space information during travel. Together, these results suggest that in CA3, but not DG, self-position information is disambiguated by temporal-order signals for both stationary platforms and dynamic traveling trajectories.

## Discussion

The present study demonstrates that substantial numbers of neurons in both CA3 and DG encode self-position while common marmosets perform the VGT task. We identified two types of position-selective neurons: platform neurons, which responded when the animals stayed at a particular platform, and trajectory neurons, which responded when the animals traveled along a specific route between platforms. Neurons exhibiting both platform- and trajectory-selective responses (conjunctive neurons) were significantly more frequent in CA3 than in DG, implying a more complex representation of spatial information in CA3. Moreover, only CA3 neurons carried temporal-order information, in particular when coupled to self-position. Together, these findings suggest a forward transformation of spatiotemporal representations along the DG–CA3 axis in the primate hippocampus (Fig. 1E).

Despite the large number of self-position-coding neurons, consistent with rodent place cells (*8*), we found almost no neurons encoding the animals’ facing location during the VGT task. Because facing location likely approximates gaze location in freely moving marmosets, this result contrasts with previous reports of spatial view cells in the primate hippocampus (*9*). The absence of spatial view coding here may be due to the visually impoverished environment of our black-walled maze in a dark room. This interpretation aligns with the “view-center background” framework, in which neurons in the ventral visual pathway encode large background images and convey them to the hippocampus via medial temporal cortical areas (e.g., perirhinal cortex) (*10, 11*). Such view-centered background signals can specify gaze location by similarity of visual backgrounds, relatively invariant to observer position (*12*), and may be a major input to spatial view cells. Visually impoverished environment in the VGT task could therefore explain the lack of spatial view cells and the relatively simple spatial codes observed here, in contrast to the rich multiplex spatial representations reported in recent studies of freely moving nonhuman primates including virtual reality environment (*14–16*).

A central finding of this study is the strong modulation of self-position-selective responses by temporal order in CA3. This temporal-order effect differentiates repeated visits to the same location across a sequence (e.g., *A*-1st vs *A*-3rd) rather than encoding the serial structure across locations within a trial (e.g., *A*→*B*→*C*) (*17, 18*). Such temporal disambiguation is a hallmark of pattern separation for similar events in both humans (*19, 20*) and animals (*6, 21*). Prior experimental and modeling work (*4, 5*) has emphasized a selective role of DG in pattern separation. However, in our task, DG spatial representations were not disambiguated by temporal order of travel, whereas CA3 integrated space and time into conjunctive spatiotemporal codes (e.g., “Platform *B* at the second temporal order”). This is in line with models proposing a strong auto-associative mechanism in CA3 (*4, 5*), capable of binding multiple dimensions such as where and when.

Assuming that CA3 constructs such spatiotemporal representations raises the question of the origin of the time signal. DG clearly provides spatial input (self-position in the maze) to CA3, but we did not detect any time-related signal independent of space in either DG or CA3 at the level of spike firing of single neurons. A comparable phenomenon has been reported for object-based temporal-order coding in the macaque medial temporal lobe, where monkeys encoded both object identity and the temporal order of two sequential stimuli (*22*). In that study, perirhinal cortex (PRC) represented integrated “what–when” information when an object was presented, without showing an explicit time signal independent of object responses, despite strong object-selective input presumably from upstream TE. In contrast, an incremental time signal during the delay period—bridging one object presentation to the next—was observed in the hippocampus, suggesting that this hippocampal time signal spread back to PRC and modulated object-selective responses to form combinatorial “what–when” codes (*22–24*). In the present study, we did not observe incremental time signals in either DG or CA3. Given rodent data showing time cells predominantly in CA1 (*7, 25*), though also present in CA3 (*26*), one possibility is that the temporal-order effects in CA3 during the VGT task are driven by a time signal originating in CA1. An alternative explanation is that our task design was suboptimal for revealing classical time cells: the VGT task neither required the animals to remain at a fixed location for a constant duration nor to traverse trajectories with fixed travel times. Future work will be needed to identify the brain regions that supply time signals to CA3 and PRC, and to elucidate how these signals are combined with space and object information to form integrated episodic representations.

## Acknowledgments

We thank Drs. L. Gao at ZIINT, X. Wang at Tsinghua University, D. Leopold at NIMH and KW. Koyano at QST for helpful discussion. We thank S. Xue for expert animal care. We thank Drs. J. Gao, W. Men, G. Yang, and the National Center for Protein Sciences at Peking University for assistance with MRI scanning. AI-assisted technologies (Google Gemini) were used for correcting grammar and clarity.

## Author contributions

Conceptualization: YN

Methodology: FQ, ARW, AN, YS, YN

Investigation: FQ

Visualization: FQ

Funding acquisition: YN

Project administration: YN

Supervision: YS, YN

Writing – original draft: FQ, YN

Writing – review & editing: FQ, ARW, AN, YS, YN

## Supplementary Materials

### Materials and Methods

#### Subjects

Two adult female common marmosets (Callithrix jacchus) (monkey B: 480 g; monkey F: 360 g) were used in all experiments. Animals were maintained on a 12-h light/dark cycle in a temperature-controlled room (26–29°C; 40–60% humidity). Health and welfare were monitored daily. Environmental enrichment and feeding followed the NIH Guide for the Care and Use of Laboratory Animals and AAALAC guidelines. All procedures were approved by the Institutional Animal Care and Use Committee of Peking University (license: Psych-YujiNaya-1).

#### Behavioral Task

The animals were trained on a visually guided travel (VGT) task in a spatial maze (2.0 m ×1.5 m ×0.7 m; Fig. 1A). The floor and walls were made of black acrylic panels covered with matte black wallpaper. Four white circular platforms (10 cm diameter, 1.0 cm height) with LEDs installed underneath were placed on the floor and designated as the Home platform (Platform *H*) and three test platforms (Platforms *A*, *B*, and *C*; Fig. 1A). A reward sipper was mounted on the wall adjacent to Platform *H*, 10 cm above the floor. Rewards (0.25 ml per trial) consisted of orange juice mixed with baby rice cereal (Gerber Products Company, USA) and were delivered by a syringe driven by a stepping motor (YK39HS56-2804A, Xiamen Yoho Automation Technology Co., Ltd., China).

The maze was covered with a transparent acrylic plate. A ceiling-mounted video camera (CinePlex, Plexon Inc., USA), positioned 2.22 m above the maze, tracked the animals’ head position by detecting two LEDs (one blue, one red) attached to the left and right sides of a wireless headstage (W32, Triangle BioSystems International, USA). The camera captured the entire maze at 30 Hz, providing real-time estimates of head position and facing direction. Position signals were transmitted to MonkeyLogic, which controlled LED illumination and reward delivery in closed loop. All sessions were conducted in a dark room.

In the VGT task, each trial began when the LED under the Platform *H* was illuminated. A “platform-zone” was defined as a circular region with a 15-cm radius centered on each platform. A trial was initiated when the marmoset’s head entered the Platform-*H*-zone and remained there continuously for 0.3 s, following that the Platform *H* LED was turned off and the LED under one of the three test platforms (Platform *A*, *B*, or *C*) was turned on. When the animal entered the cued platform-zone and stayed there for 0.3 s, the LED on that platform turned off. The animal was then required to remain within the same platform-zone for an additional 1.0 s, which triggered illumination of the next test platform (Fig. 1B). The animal was required to move to the illuminated test platform. After the animal visited the 3rd test platform and stayed there for 1.3 s in total, a reward (0.25 ml) was delivered only. Trials were terminated without reward if the animal failed to reach the cued platform or to remain within its platform-zone for a total of 1.3 s. Six possible sequences (*A*–*B*–*C*, *A*–*C*–*B*, *B*–*A*–*C*, *B*–*C*–*A*, *C*–*A*–*B*, *C*–*B*–*A*) were presented pseudo-randomly within each session (Fig. 1C). Sessions with at least 24 completed trials (24 visits to each test platform) were included in subsequent analyses.

#### Electrophysiological Recording

For surgery, animals were sedated with intramuscular atropine (0.05 mg/kg, i.m.) and Zoletil 50 (one to one mixture of tiletamine hydrochloride and zolazepam hydrochloride, 15 mg/kg, i.m.), placed in a stereotaxic frame (Narishige, Japan), and maintained under 1.0–2.0% isoflurane anesthesia. Heart rate, body temperature, and arterial oxygen saturation (SpO₂) were monitored (Cardell Touch, Midmark Corporation, USA) and maintained at >200 beats/min, >36°C, and >90%, respectively. The scalp was incised, and the skull was widely exposed. The exposed skull was decalcified with 10% citric acid (1 min) followed by 0.3% hypochlorous acid (1 min), then rinsed with saline. The decalcified surface was coated with a dental adhesive resin (PANAVIA V5, Kuraray, Japan) and overlaid with dental resin (Unifast II, GC Corporation, Tokyo, Japan), forming a stable base (*27*). A resin recording chamber was then affixed to the skull, and a connector (Plexon Inc., USA) was attached to the anterior portion, leaving access for subsequent electrode implantation (Fig. S1B).

Using individual anatomical MRI acquired preoperatively on a 9.4 T scanner (Bruker, Germany), 16-channel microwire brush array (MBA) electrodes (Microprobes, Gaithersburg, USA) mounted on a microdrive were chronically implanted into the left hippocampus (monkey B: 3.3 mm anterior to the external auditory meatus, 7.5 mm lateral from the midline; monkey F: 4.8 mm anterior, 7.2 mm lateral), adapting published approaches (*28,29*). Electrode positions were monitored physiologically during implantation and verified postsurgically by CT imaging co-registered to the preoperative MRI (Fig. 1D and Fig. S1C). The MBA arrays were advanced with a 300-µm pitch drive screw (*30,31*), allowing fine post-implantation adjustment. During recording, the microdrive was lowered in 37.5–75.0 µm increments per day to sample neurons across hippocampal subregions.

Neural signals were bandpass-filtered for single units (200 Hz–6 kHz), digitized at 40 kHz (OmniPlex, Plexon), and stored for offline analysis. After completion of recordings, hippocampal subregions in monkey B were identified histologically using Nissl staining (Fig. 1D). The dentate gyrus (DG) was characterized by a densely packed, curved granule cell layer, bordered by a sparsely cellular molecular layer and a heterogeneous polymorphic layer. The CA3 showed a compact pyramidal cell layer of large triangular neurons with strong Nissl staining, with apical and basal dendrites projecting to the molecular layer and stratum oriens, respectively.

Monkey F is still being used; thus, histological verification is pending. For this animal, recording sites were instead localized by CT–MRI co-registration (*32*) combined with physiological criteria. The discrimination of the CA3 and DG subregions was confirmed using the electrophysiological data. The CA3 subregion is located in the superficial layer while the dentate gyrus (DG) is in the deep layer, with each layer spanning approximately 400 μm. A spike-free gap exists between CA3 and DG; thus, the specific hippocampal subregion can be accurately identified based on the presence and absence of spikes.

#### Data Analysis

High-pass neural signals were spike-sorted offline using the OmniPlex Offline Sorter (Plexon Inc., USA) with manual curation to ensure that noise transients were not included as units and that single cells were not split into multiple clusters. Because noise was unavoidable in freely moving recordings, data from a given platform or trajectory condition were discarded if noise contaminated >5% of either the total time spent in the platform zone or the total trajectory duration. Spikes occurring during sharp-wave ripple (SWR) events when animals were in a platform zone were excluded (*33*). SWRs were defined as periods in which ripple-band (100–250 Hz) power exceeded the mean by >2 s.d. Processed data were analyzed in MATLAB (MathWorks) using custom scripts and the Statistics and Machine Learning Toolbox.

##### Platform selectivity

To quantify platform-selective responses, we computed the mean firing rate for each neuron during the period when the animal stayed at each test platform. For each trial, this analysis window started 300 ms after entry into the platform-zone and ended when the animal left the platform-zone for the next LED-illuminated target platform or for reward collection. If the animal continued to stay in the platform-zone for >1.5 s after the next test platform-LED illumination or reward delivery, data were included up to a maximum total analysis window of 2.8 s (1.3 s + 1.5 s). For each neuron, we then performed a one-way ANOVA with platform (three levels: *A*, *B*, *C*) as the main factor. Neurons with significantly different firing rates across platforms (*P* < 0.01) were defined as platform neurons.

##### Trajectory selectivity

To quantify trajectory-selective responses, we computed the mean firing rate for each neuron during each traversal from one platform-zone to another (e.g., *A*→*B*) on each trial. The ideal straight-line distance between platform-zones was 0.85 m (1.15 m − 2 × 0.15 m). For each neuron, we performed a one-way ANOVA with trajectory (six levels: *A*→*B*, *A*→*C*, *B*→*A*, *B*→*C*, *C*→*A*, *C*→*B*) as the main factor. Neurons with significantly different firing rates across trajectories (*P* < 0.01) were defined as trajectory neurons. Neurons that were either platform neurons or trajectory neurons were considered to be “position” neurons.

##### Temporal-order effects

Temporal-order effects were examined separately for platform and trajectory responses. For platforms, we applied a two-way ANOVA to each neuron with factors platform (three levels: *A*, *B*, *C*) and temporal order (three levels: 1st, 2nd, 3rd). For trajectories, we used a two-way ANOVA with trajectory (six levels) and temporal order (two levels) as main factors. For both analyses, we evaluated main effects of temporal order and time ×space (platform or trajectory) interactions, using a significance threshold of *P* < 0.01. Neurons showing either a significant time main effect or a significant interaction were classified as temporal-order neurons.

##### Facing location and directional tuning

To test whether platform responses could be explained by facing location, we projected the animal’s facing direction at each test platform (*A*, *B*, *C*) onto the surrounding maze walls and computed mean firing rates in 50-cm bins along the projected walls across trials, yielding a 14-bin firing-rate vector for each platform and neuron. For platform neurons, we quantified the similarity between the firing-rate vector at the best-preferred platform and those at the other platforms by computing two Pearson correlation coefficients (preferred vs. each non-preferred platform) and averaging them. For non-platform neurons, we computed three Pearson correlation coefficients for all platform pairs (*A*–*B*, *A*–*C*, *B*–*C*) and averaged them. Bins with a total occupancy <1 s were excluded. We also computed Pearson correlation coefficients for firing rates between trajectory pairs with the same goal (*AB* vs. *CB*, *BA* vs. *CA*, *AC* vs. *BC*), as well as between trajectory pairs with the same physical position but different goals (*AB* vs. *BA*, *AC* vs. *CA*, *BC* vs. *CB*). If no spikes occurred in any bin of a trajectory or facing-direction vector (i.e., an all-zero vector for which Pearson’s r is undefined), we assigned r = 0.0 for subsequent analyses. In addition to Pearson’s r, we applied non-parametric tests to assess the distributions of correlation values (Fig. 2D, S3C, and S4F).

Directional tuning functions were constructed by plotting firing rate as a function of the marmoset’s facing direction, binned into 10° intervals and smoothed with a Gaussian kernel (one-bin smoothing on each side). Directional tuning strength was quantified by the mean vector length (MVL) of the circular firing-rate distribution across all direction bins. A neuron was classified as significantly direction-tuned if its MVL exceeded the 99th percentile of MVL values obtained from shuffled permutations of its original firing-rate data.

## Supplementary Text

### Tuning properties of platform neurons and trajectory neurons

We examined tuning properties of platform neurons in both regions. Across neurons, the strongest responses were observed not only for Platform *B*, as in the example neuron (Fig. 2A), but also for Platforms *A* and *C* (Fig. S2B). Spatial tunings of platform neurons were characterized by the ratios of firing rates at the middle- and least-preferred platforms relative to the best platform (Fig. S2C; median values: 0.53, 0.22 for CA3; 0.38, 0.14 for DG). Michelson contrasts of position-selective activity did not differ between CA3 and DG (*P* = 0.610, *KS* = 0.141, Kolmogorov Smirnov test Fig. S2D). Together with the results presented in Fig. 2B and Table 1, the two hippocampal subfields carried substantial self-position signal when the animals stayed at the test platforms although self-position signals were slightly stronger in CA3 than DG (Fig. S2A).

Trajectory neurons encoded their best-preferred trajectory almost exclusively in both hippocampal subfields, as indicated by the response ratio of the second-best trajectory to the best trajectory (median = 0.17 and 0.23 for CA3 and DG, respectively; Fig. S4B, also see Fig. S4C), while the best-preferred trajectories collectively covered all six trajectories (Fig. S4B). The normalized response difference between the best and second-best trajectories did not differ between CA3 and DG (*P* = 0.94, *KS* = 0.13; Fig. S4E).

## Supplementary figures

**Fig. S1.**
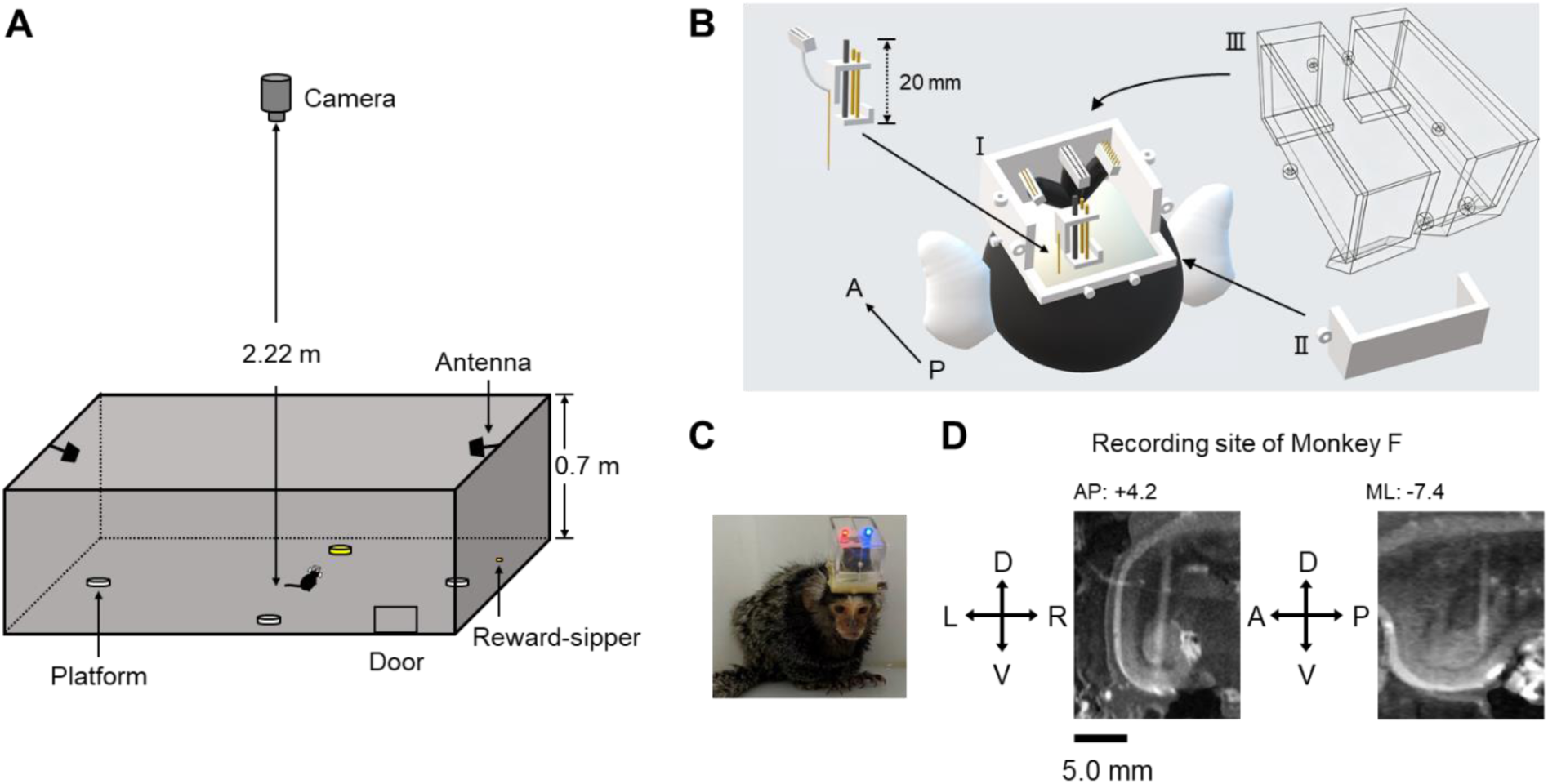
Experimental setup and apparatus. (A) 3D schematic of the behavioral maze. (B) Schematic of the chronic recording chamber (Ⅰ), detachable rear wall (Ⅱ) and protective transparent lid (Ⅲ), with microelectrodes mounted on the microdrive. (C) Photograph of Monkey F with implanted headstage. (D) Combined MRI and CT images showing the electrode location relative to brain structure of Monkey F.

**Fig. S2.**
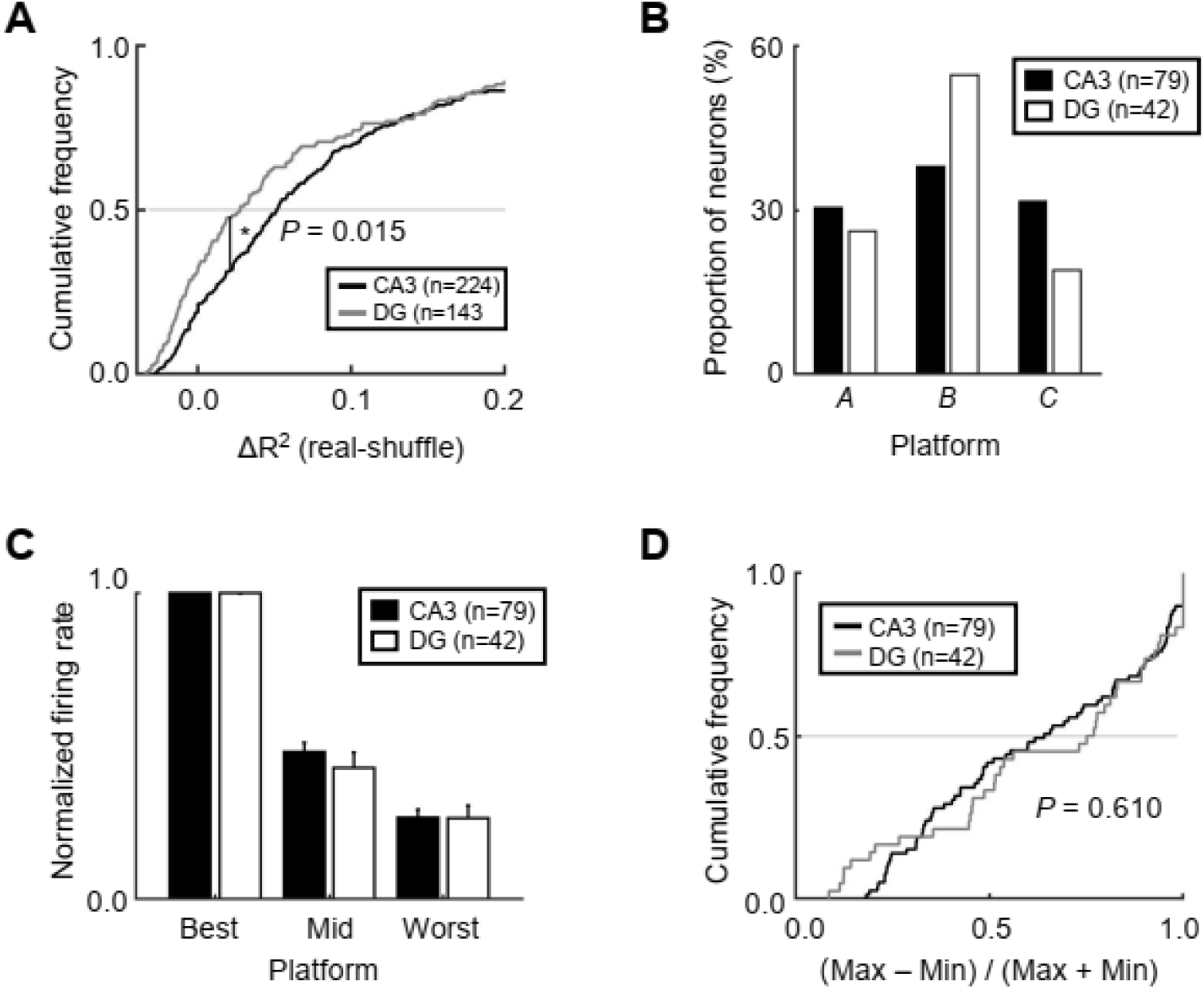
Platform selective activities. (A) Cumulative frequencies of recorded neurons were plotted against *R*^2^ values reflecting their response selectivity at staying test platforms in CA3 and DG. The *R*²value was subtracted by a mean of 1,000 shuffled *R*²values for each neuron. The subtracted *R*²values were significantly larger in CA3 than in DG (*P* = 0.015, *KS* = 0.166; Kolmogorov–Smirnov test). (B) Proportions of platform neurons by their best-preferred platforms. (C) Tuning property of self-position selective activity among three test platforms. The best-preferred platform, middle-preferred platform and worst-preferred platform were determined for each platform neuron, and then firing rate at each test platform was normalized by the firing rate at the best-preferred platforms. Bar graphs indicate means and SEMs of normalized firing rates for platform neurons. (D) Cumulative frequencies of platform neurons were plotted against the contrast of their platform-selective responses in each area. The two subfields did not show a significant difference in the contrast (*P* = 0.61, *KS* = 0.14; Kolmogorov–Smirnov test).

**Fig. S3.**
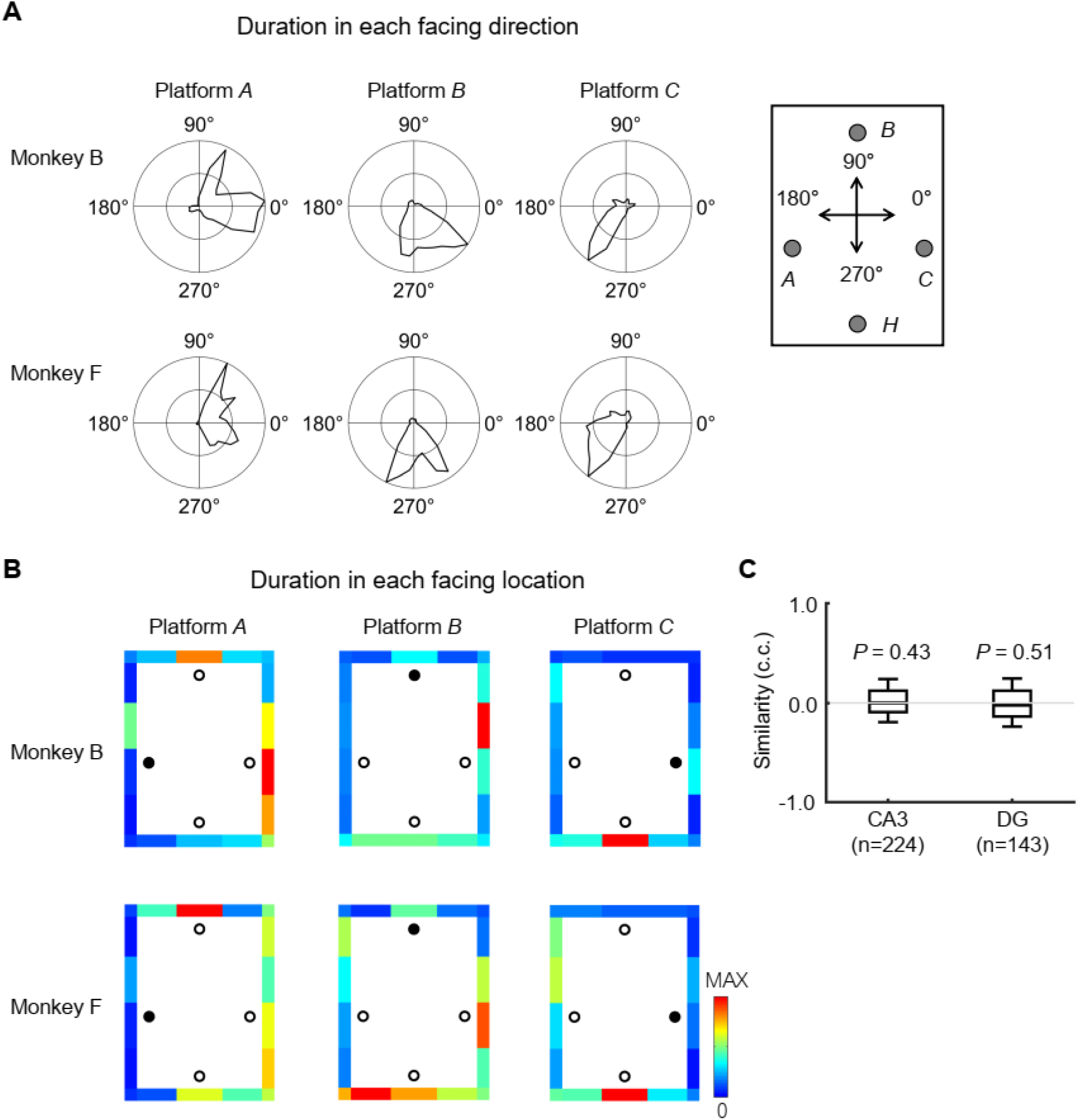
Firing pattern similarities for facing locations between platforms. (A) Time duration distributions of marmosets’ facing directions from different test platforms. (B) Time distributions of marmosets’ facing locations from different test platforms. The circle represents the platform, with black filled circles indicating marmoset positions. (C) Firing pattern similarities for facing locations were examined between all pairs of test platforms. For each neuron, the mean pattern similarity across the three pairs (Platform *A*-Platform *B*, Platform *B*-Platform *C* and Platform *C*-Platform *A*) was calculated. Box-and-whisker plots indicate distributions of the mean firing pattern similarities for all recorded neurons in CA3 and DG. The horizontal line within each box represents the median, while the upper and lower boundaries of the box indicate the interquartile range. The whiskers extend to the 10th and 90th percentiles, representing the extreme data points excluding outliers.

**Fig. S4.**
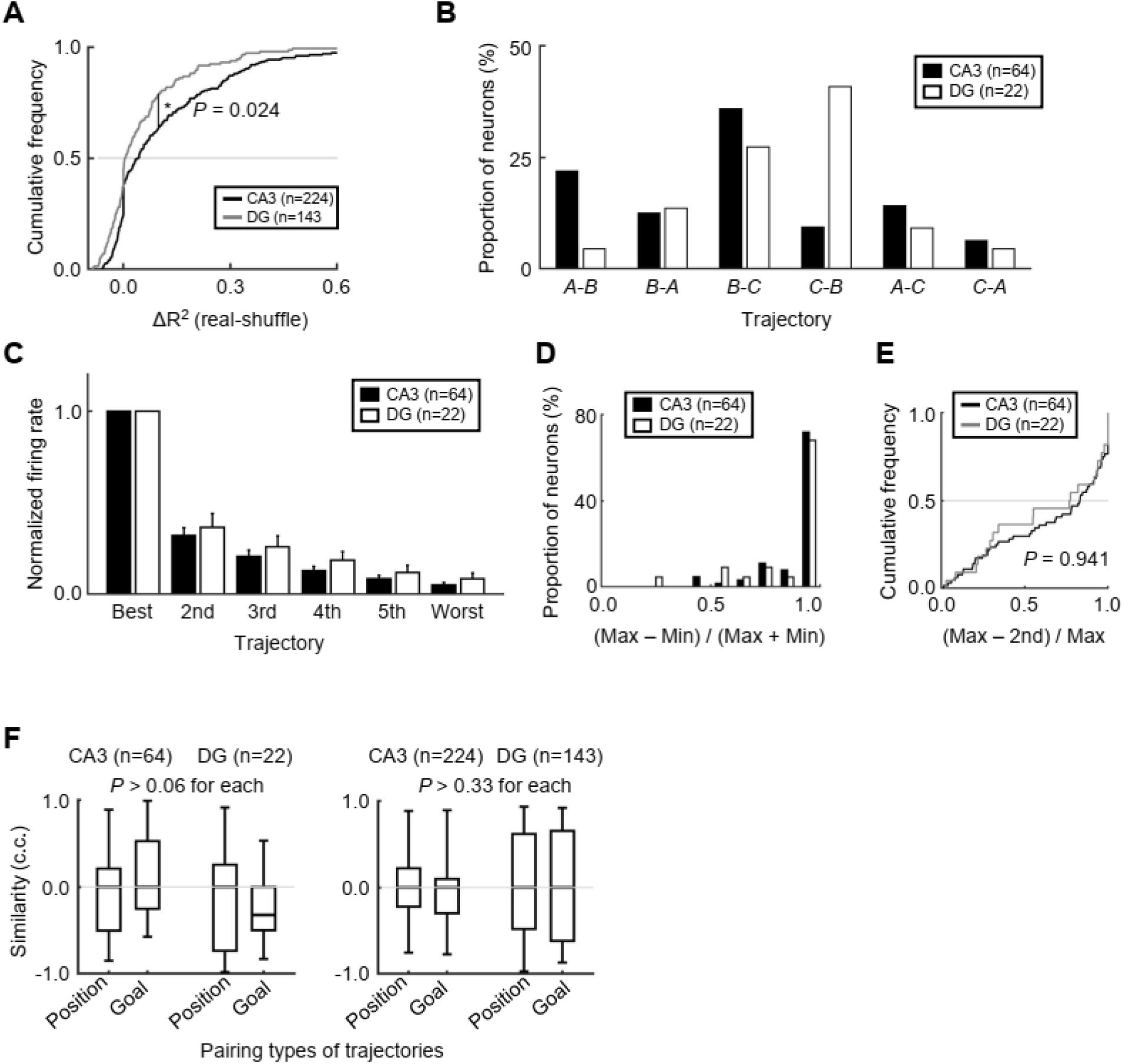
Representation of trajectory selective activities. (A) Cumulative frequencies of recorded neurons were plotted against *R*^2^ values reflecting their response selectivity to trajectories during traveling from one test platform to another in CA3 and DG. The *R*²value was normalized by subtracting a mean of 1,000 shuffled *R*²values for each neuron. The normalized *R*²values were significantly larger in CA3 than in DG (*P* = 0.024, *KS* = 0.16; Kolmogorov–Smirnov test). (B) Proportions of trajectory neurons categorized by their best-preferred trajectories. (C) Tuning property of self-position selective activity across six trajectories. For each trajectory neuron, the best-preferred platform, 2nd-preferred and worst-preferred trajectories were determined. A firing rate along each trajectory was normalized by the firing rate along the best-preferred trajectory. Bar graphs indicate mean ±SEM of normalized firing rates for trajectory neurons. (D) Contrast for trajectory neurons, defined as the normalized difference between the maximum and minimum mean firing rates across the six trajectories. (E) Cumulative number of trajectory neurons plotted against their response tunings calculated as the difference between the maximum and second-highest mean firing rates relative to the maximum firing rate (*P* = 0.941, *KS* = 0.126; Kolmogorov–Smirnov test). (F) Firing pattern similarities were examined between trajectories with same positions but opposite directions (Position) and with different positions but same goal platforms (Goal). Box-and-whisker plots indicate distributions of the mean firing pattern similarities for trajectory neurons (left) and recorded neurons (right) in CA3 and DG. The horizontal line within each box represents the median, while the upper and lower boundaries of the box indicate the interquartile range. The whiskers extend to the 10th and 90th percentiles, representing the extreme data points excluding outliers.

**Fig. S5.**
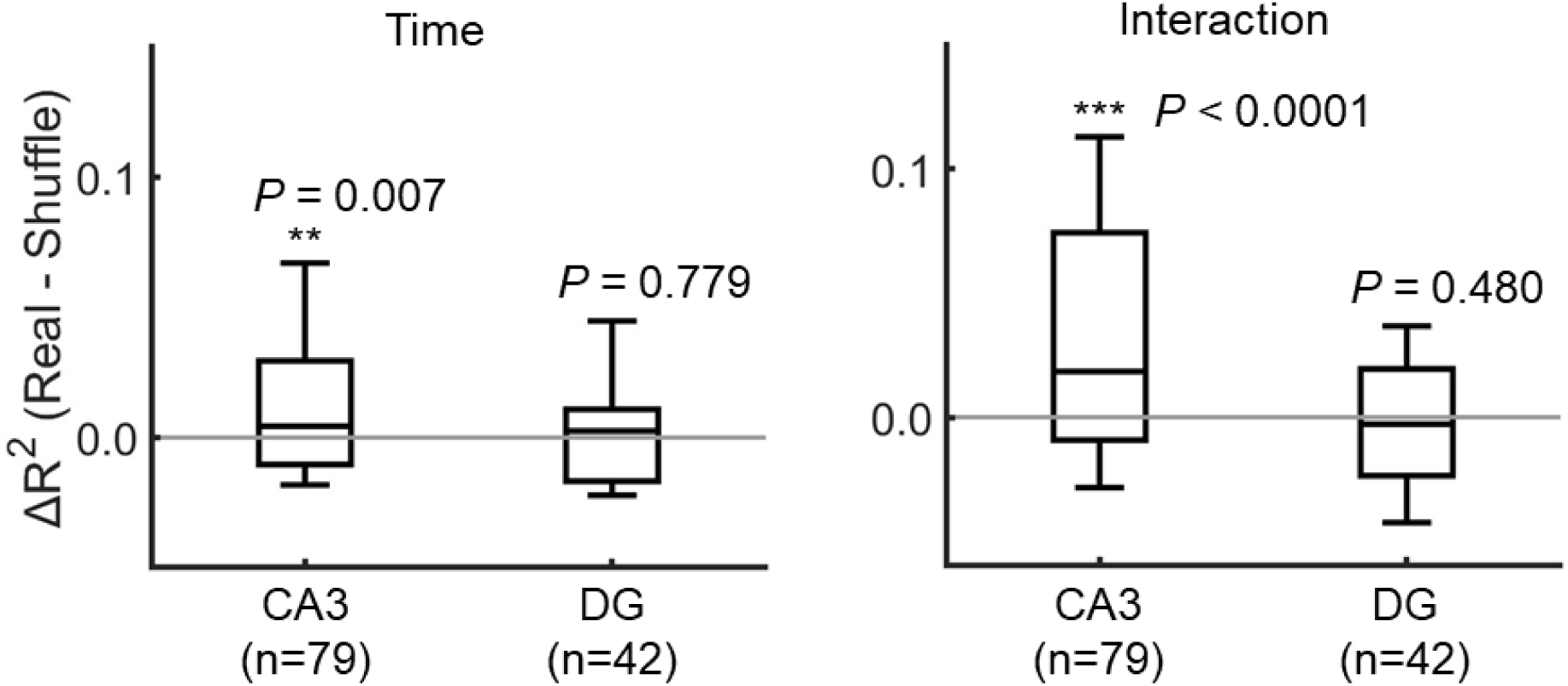
Temporal-order effect of CA3 and DG. Box-and-whisker plots indicate the distributions of *R*²values reflecting the main effect of time (left) and space ×time interaction (right) for platform neurons in CA3 and DG. The horizontal line within each box represents the median, while the upper and lower boundaries of the box indicate the interquartile range. The whiskers extend to the 10th and 90th percentiles, representing the extreme data points excluding outliers. For each neuron, the *R*²value was normalized by subtracting the mean of 1,000 shuffled *R²*values. Main effect of time: CA3: ** *P* = 0.007, *Z* = 2.69; DG, *P* = 0.779, *Z* = 0.281. Space × time interaction: CA3: *** *P* < 0.0001, *Z* = 4.24; DG, *P* = 0.48, *Z* = -0.707 (Wilcoxon signed-rank test, two-tailed).

**Fig. S6.**
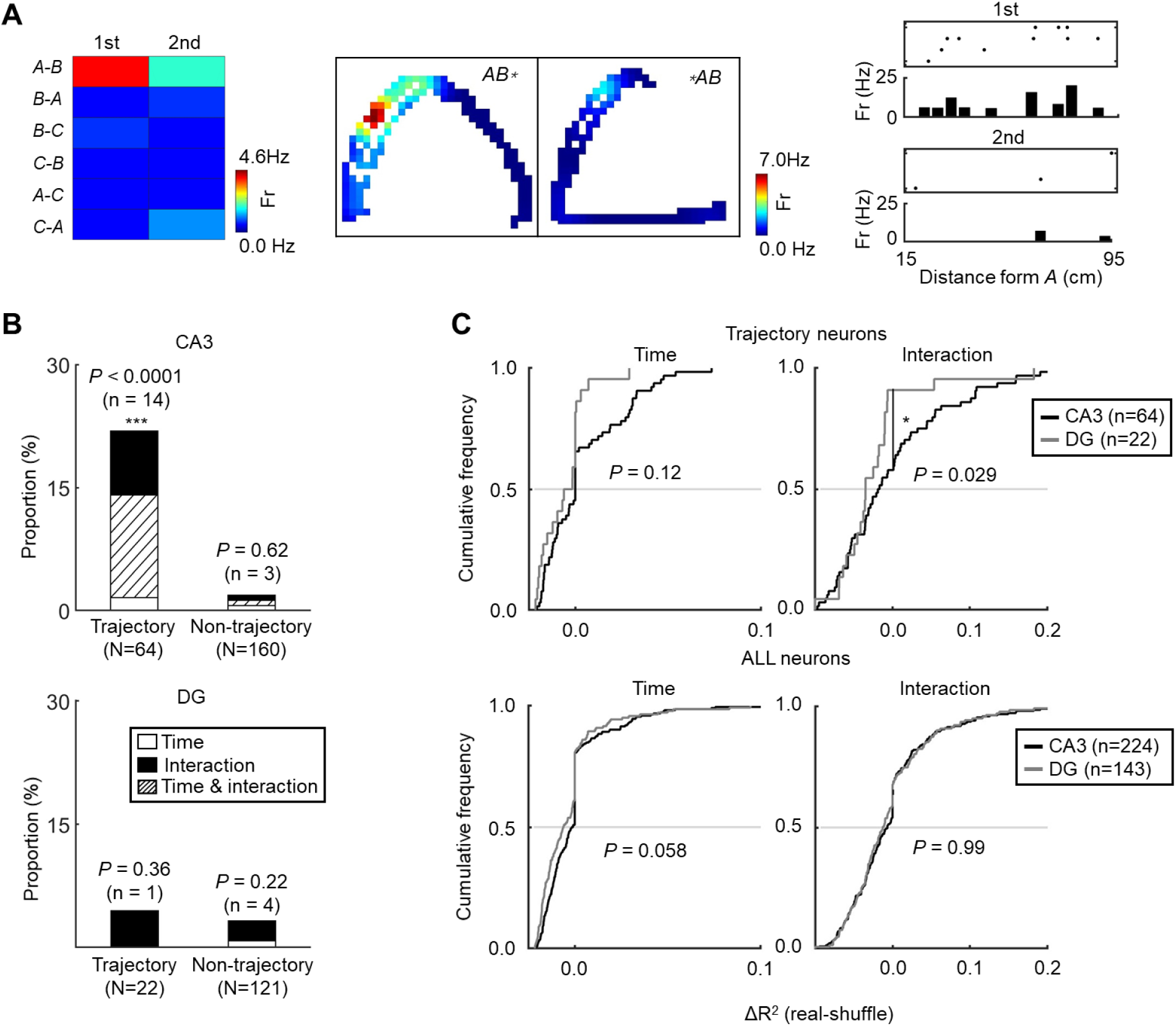
Representation of temporal-order effects in trajectory neurons. (A) A representative CA3 neuron showing the temporal-order effect on trajectory selective activities. Firing rate maps and raster plots indicate that neuronal firing was maximal during the first temporal order of the animal’s trajectory from platform *A* to *B* (*A*-*B*). (B) Bar graphs showing the proportions of temporal-order neurons, categorized by the main effect of time and the interaction effect, among trajectory neurons (left) and non-trajectory neurons (right). Number of temporal neurons among trajectory neurons was significantly larger than the chance level (2%) on CA3 (*** *P* < 0.0001, *Z* = 11.4, binominal test). (C) Cumulative frequency of CA3-DG differences in the main effect of time and interaction effects for trajectory neurons (top) and all recorded neurons (bottom), quantified as the *R*^2^ values normalized by subtracting the mean of 1,000 shuffled *R²*values (\**P* = 0.029, Kolmogorov–Smirnov test).

## Supplementary Tables

**Table. S1.** Duration of staying at test-platforms and traveling between test-platforms.

|  | **25 %** | **Median** | **75 %** |
| --- | --- | --- | --- |
| **Staying at platforms** |  |  |  |
| **1st** | 2.37 | 2.98 | 4.53 |
| **2nd** | 2.37 | 3.10 | 4.53 |
| **3rd** | 2.13 | 2.57 | 3.63 |
| **Traveling between platforms** |  |  |  |
| **1st** | 0.53 | 0.57 | 0.63 |
| **2nd** | 0.53 | 0.57 | 0.60 |

5 The scale of the duration is in seconds. Data were obtained from 1,896 trials across 62 recording sessions (Monkey B: 1,141 trials / 35 sessions; Monkey F: 755 trials / 27 sessions).

**Table S2.** Numbers of neurons encoding the temporal order of visited platforms.

|  | **Recorded** | **Temporal-**<br>**order** | **Main (time)** | **Interaction**<br>**(time× space)** |
| --- | --- | --- | --- | --- |
| **Total** |  |  |  |  |
| **CA3** | 224 | 44 (19.6%) | 25 (11.2 %) | 31 (13.8 %) |
| **DG** | 143 | 3 (2.1 %) | 3 (2.1 %) | 1 (0.7 %) |
| **Monkey B** |  |  |  |  |
| **CA3** | 151 | 28 (18.5 %) | 16 (10.6 %) | 20 (13.3 %) |
| **DG** | 46 | 1 (2.2 %) | 1 (2.2 %) | 1 (2.2 %) |
| **Monkey F** |  |  |  |  |
| **CA3** | 73 | 16 (21.9 %) | 9 (12.3 %) | 11 (15.1 %) |
| **DG** | 97 | 2 (2.1 %) | 2 (2.1 %) | 0 (0.0 %) |

Temporal-order neurons were defined as those exhibiting a significant main effect of time or a significant time × space interaction ( $P < 0.01$ for each, two-way ANOVA). Percentages in parentheses indicate the proportions of significant neurons relative to the total number of recorded neurons.

**Table S3.** Numbers of neurons encoding the temporal order of travelled trajectories.

|  | **Recorded** | **Temporal-order** | **Main (time)** | **Interaction (time×space)** |
| --- | --- | --- | --- | --- |
| **Total** |  |  |  |  |
| CA3 | 224 | 17 (7.6 %) | 11 (4.9 %) | 15 (6.7 %) |
| DG | 143 | 5 (3.5 %) | 1 (0.7 %) | 4 (2.8 %) |
| **Monkey B** |  |  |  |  |
| CA3 | 151 | 13 (8.6 %) | 9 (6.0 %) | 11 (7.3 %) |
| DG | 46 | 1 (2.2 %) | 0 (0 %) | 1 (2.2 %) |
| **Monkey F** |  |  |  |  |
| CA3 | 73 | 4 (5.5 %) | 2 (2.7 %) | 4 (5.5 %) |
| DG | 97 | 4 (4.1 %) | 1 (1.0 %) | 3 (3.1 %) |

Temporal-order neurons were defined as those exhibiting a significant main effect of time or a significant interaction ( $P < 0.01$ for each, two-way ANOVA). Percentages in parentheses indicate the proportions of significant neurons among the total number of recorded neurons.

## Supplementary movies

**Movie S1.** Behavioral video of a marmoset traversing the platform sequence *A* → *B* → *C* during one trial.

**Movie S2.** Behavioral video of a marmoset traversing the platform sequence *B* → *A* → *C* during one trial.

**Movie S3.** Behavioral video of a marmoset traversing the platform sequence *B* → *C* → *A* during one trial.

